# Restricted access to beneficial mutations slows adaptation and biases fixed mutations in diploids

**DOI:** 10.1101/171462

**Authors:** Daniel A. Marad, Gregory I. Lang

**Affiliations:** Department of Biological Sciences, Lehigh University, Bethlehem PA 18015

## Abstract

Ploidy varies considerably in nature. Yet, our understanding of the impact of ploidy on adaptation is incomplete. Many microbial evolution experiments characterize adaptation in haploid organisms, but few focus on diploid organisms. Here, we perform a 4,000-generation evolution experiment using diploid strains of the yeast *Saccharomyces cerevisiae*. We show that the rate of adaptation and spectrum of beneficial mutations are influenced by ploidy. Haldane’s sieve effectively restricts access to beneficial mutations in diploid populations, leading to a slower rate of adaptation and a spectrum of beneficial mutations shifted towards dominant mutations. Genomic position also plays an important role, as the prevalence of homozygous mutations is largely dependent on their proximity to a recombination hotspot. Our results demonstrate key aspects of diploid adaptation that have previously been understudied and provide support for several proposed theories.

## INTRODUCTION

Understanding the impact of ploidy on adaptation is a central challenge in evolutionary biology. Ploidy varies considerably in the natural world from bacteria that are mostly haploid to some plants that can exist as decaploid^1^. In addition, all sexual organisms alternate between ploidy states through gamete fusion and meiosis^2^. Despite its importance, we have an incomplete picture of how ploidy impacts the rate of adaptation and spectrum of beneficial mutations during adaptive evolution. In principle, how ploidy impacts adaptation depends largely on assumptions regarding the dominance of new beneficial mutations. If new beneficial mutations are mostly dominant, then diploids, with twice the mutational target size as haploids, will be twice as likely to acquire beneficial mutations, and will have greater evolutionary potential. Alternatively, if beneficial mutations are recessive, then haploids will have access to new beneficial mutations that have no selective advantage in diploid populations^3^. This filtering of recessive beneficial mutations in diploids was first proposed by Haldane^4^ and thus is commonly known as Haldane’s sieve. It is likely that Haldane’s sieve is not the only factor affecting the impact of ploidy on adaptation and that other factors, such as deleterious load and physiological differences between haploids and diploids, also contribute to differences in adaptive potential. If deleterious mutations are mostly recessive, negative phenotypes will be largely masked in diploids. However, if deleterious mutations are dominant (e.g. haploinsufficient mutations), diploids would be negatively impacted by having twice the deleterious load as haploids. In addition, it has been shown in yeast that different ploidy states experience differential regulation of some genes^5^, which may further impact the mutations that are selectively accessible in haploids and diploids.

The budding yeast, *Saccharomyces cerevisiae*, can be stably propagated asexually in both haploid and diploid states. This provides an ideal system for studying the effect of ploidy on adaptation. Early experimental results using yeast seemed to demonstrate that diploids evolved faster^6^. Paquin and Adams looked at rates of adaptation indirectly by monitoring the purging of neutral markers as a proxy for the frequency of selective sweeps and found that diploids fixed mutations 60% faster than haploids, and that this rate did not slow over time^6^. However, despite this early work, most recent studies find that haploid yeast adapt more quickly compared to diploids, and that this result holds across many strain backgrounds and many environments^7–10^. Yet, why haploids adapt more quickly is unclear.

Extensive sequencing of laboratory-evolved populations has focused almost exclusively on the spectrum of beneficial mutations in bacterial and haploid yeast populations, rather than in diploids^11–14^. Collectively, this work shows that the spectrum of beneficial mutations in haploid populations is skewed towards loss-of-function mutations. While there is less information regarding the spectrum of beneficial mutations in higher ploidy populations, one recent effort used whole-genome sequencing and gene expression analysis to explore adaptation of polyploid yeast (including diploids) to Raffinose media and observed a broader spectrum of beneficial mutations in tetraploids^15^. Some work in diploids implicates Haldane’s sieve as a filter for recessive beneficial mutations. For example, Gerstein *et al*. constructed heterozygous diploids from a collection of nystatin-resistant haploid yeast, and found that all evolved nystatin-resistant mutations were recessive^16^. Recent theory also suggests that beneficial mutations in diploids may be overdominant (more beneficial as heterozygotes than homozygotes)^17^ and there is some experimental evidence suggesting this is the case^10,18^. Based on these predictions, there is a need to determine how diploidy changes the spectrum of beneficial mutations and the dynamics of adaptation.

Here we measure the rate of adaptation for 48 replicate diploid populations through 4,000 generations and compare these results to previously evolved haploid populations^13^. We sequence two clones each from 24 populations after 2,000 generations. For two populations we perform whole-genome whole-population time-course sequencing to determine the dynamics of adaptation in diploids and the effect of diploidy on the spectrum of fixed heterozygous and homozygous mutations. We show that diploids adapt more slowly than haploids, have a unique spectrum of beneficial mutations, and that the prevalence of homozygous mutations depends on their genomic position. In addition, we validate haploid-specific, diploid-specific, and shared mutational targets by reconstruction.

## RESULTS

### Most of the fitness advantage of mutations in evolved haploids is recessive

Previously, we determined the spectrum of mutations in haploid populations evolved for 1,000 generations^13^. We observe a large number of nonsense and frameshift mutations, suggesting that adaptation in haploids may be driven largely by loss-of-function mutations. Many of these mutations are likely to be recessive given that only 3% of gene deletions are haploinsufficient or haploproficient^19^. To test this hypothesis, we isolated twelve individual clones containing diverse mutations from ten evolved haploid populations and crossed them to the *MAT*α version of the ancestor to generate diploids heterozygous for all of the evolved mutations. To control for effects of mating type, the heterozygous diploids were converted to *MAT***a**/**a**. Fitness of the haploid evolved strains ranged from 2% to 8% (**Fig. 1**). Heterozygous diploid fitness, however, was only significantly different from zero in one of twelve populations. Interestingly, this one haploid population with partially dominant fitness effects contains epistatically-interacting alleles of *KEL1* and *HSL7*, large effect driver mutations that were not common targets of selection^20^. Outside of this one example, beneficial mutations in haploids are recessive. Based on this result, we expect Haldane’s sieve to filter out most of these mutations were they to occur in a diploid background, leading to a slower rate of adaptation in diploids and a spectrum of beneficial mutations that is shifted towards dominant gain-of-function mutations.

**Fig. 1:**
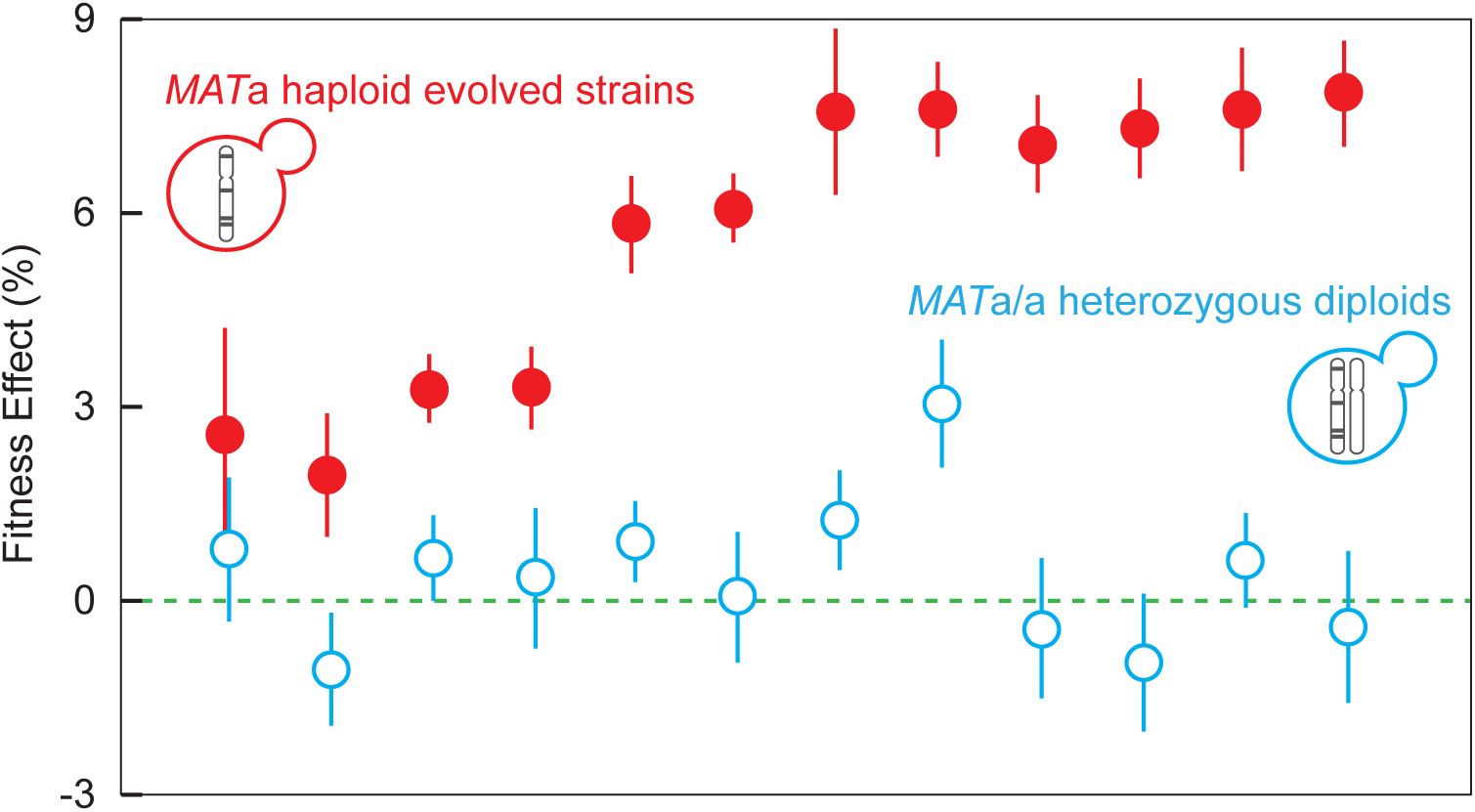
Beneficial mutations in evolved haploids are recessive. Filled red circles indicate fitness of twelve evolved *MAT***a** clones from the haploid experiment^14^. Open blue circles indicate the fitness of the same twelve strains as *MAT***a**/**a** diploids where all of the evolved mutations are heterozygous. Error bars are standard error between 7 replicates. Haploid fitness data are from Ref. 20. In order from left to right, the data points correspond to populations BYS1-E03-745, RMS1-G02-545, BYS2-C06-1000, BYS1-A08-545, BYS2-D06-910, BYS1-A09-1000, BYS-D08-1000, BYS2-E01-745, RMS1-G02-825, RMS1-H08-585, RMS1-H08-585, and RMS1-D12-910 in Refs. 13,20.

### Diploids evolve more slowly than haploids

To test our hypothesis that the rate of adaptation and mutational spectrum differs between ploidies, we evolved 48 diploid populations in rich glucose medium (YPD) for 4,000 generations under conditions identical to those used for the haploid experiment^13^. The fitness of each population was measured approximately every 250 generations using a competitive growth rate assay. On average, diploid populations increased in fitness by only 5.8% over 4,000 generations, whereas haploid populations increased in fitness by 8.5% over 1,000 generations (**Fig. 2**). As expected given the stochastic nature of evolution, we observe a large variation in fitness gains and trajectories across replicate populations. Diploids increase in fitness by between 2% and 11% over 4,000 generations, whereas haploids increase in fitness between 5% and 12% over 1,000 generations (**Fig. 2**). In order to predict future rates of adaptation, we fit the averaged data to both a power law (which has been shown to accurately model haploid evolution^21^) and a linear fit. The haploid data are described better by a power law fit than by a linear fit (*p*<0.001, F-test) but the diploid data are not (*p*=1, F-test).

**Fig. 2:**
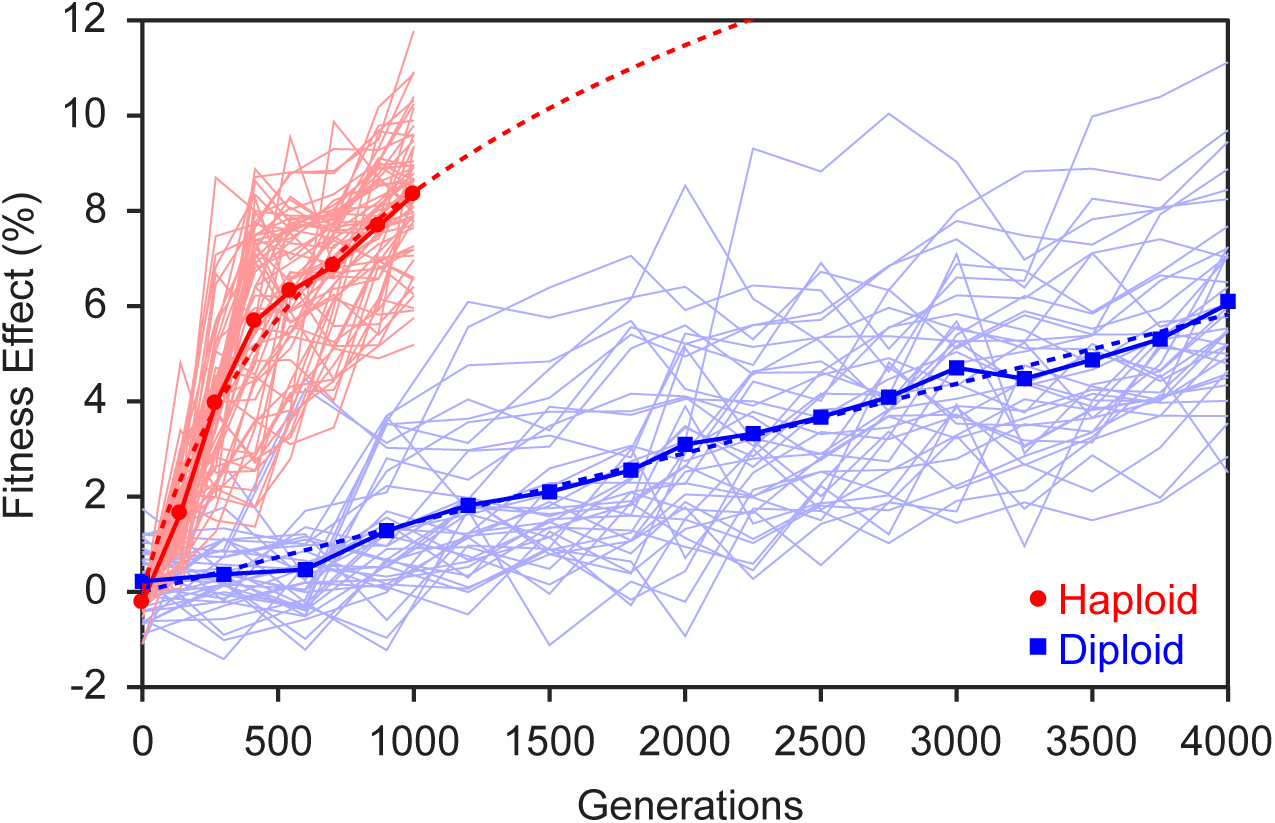
The rate of adaptation of haploid and diploid populations. Over the course of a 4,000 generation evolution experiment, diploids adapt more slowly than haploids. Fitness effects for haploids and diploids are plotted as averages represented by red circles and blue squares, respectively. Light red and light blue traces indicate individual trajectories of haploid and diploid populations, respectively. The red dotted line represent the fit of the haploid average to the power law equation, *y* = (*bx* + 1)^*a*^ −1. The blue dotted line represents the linear fit of the diploid average data. Haploid data are from Ref. 13.

### The spectrum of beneficial mutations is different between diploids and haploids

Having confirmed that diploids adapt more slowly than haploids, we next determined the spectrum of mutations in diploids by sequencing two clones each from 24 diploid populations from generation 2,000. Each clone was sequenced to an average depth of ~30x coverage and we called mutations only if they were present in both clones. We classified each mutation as heterozygous or homozygous. We identify 381 mutations across the 24 populations. Heterozygous mutations outnumber homozygous mutations 9:1. The number of mutations per population ranged from 5 in population B04 to 27 in population G08. Of the 381 mutations, 118 are intergenic, 56 are synonymous, 175 are missense, and 31 are nonsense or frameshift (**Table S1**). Comparing this to the set of haploid mutations, we see that diploids are slightly enriched for intergenic and synonymous mutations but this is not significant by Chi-square analysis (295 intergenic, 101 synonymous, 492 missense, 118 nonsense/frameshift, p=0.025). Homozygous mutations are overrepresented on the right arm of Chr. XII (*p*<0.001, Chi-square). Excluding Chr. XII, the genome-wide distribution of homozygous and heterozygous mutations is not significantly different from random (*p*=0.06, Chi-square, **Fig. 3**).

**Fig. 3:**
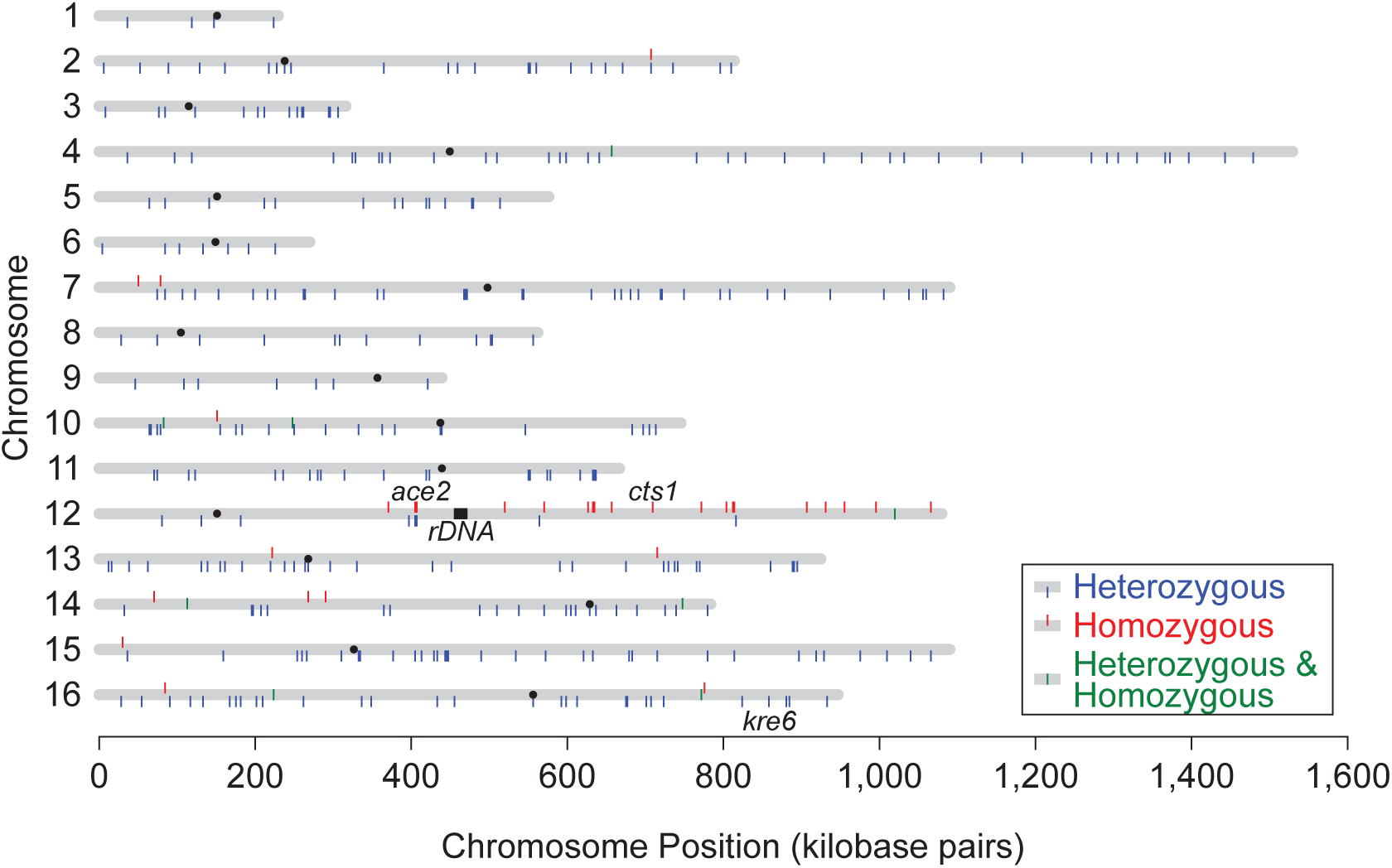
Genome-wide distribution of mutations in diploids evolved for 2,000 generations. Sequencing data for 381 diploid mutations show an overrepresentation of homozygous mutations on the right arm of Chr. XII (*p*<0.001, Chi sq.). Chromosomes are represented as grey bars. Homozygous mutations are represented by red lines sticking out of the top of chromosomes. Heterozygous mutations are represented by blue lines sticking out of the bottom of chromosomes. Mutations that were seen as both homozygous and heterozygous in paired samples are shown as green lines in the middle of chromosomes.

Of the mutations in our diploid populations, we identified putative targets of selection as those genes in which mutations were seen at least twice (**Table 1**). We observe interesting differences in the biological processes that are common targets of selection in diploids. Genes involved in the Ras pathway and the mating pathway are common targets of selection in haploid populations^13^. However, no mutations in either of these two pathways were detected in the evolved diploids. Cell wall biogenesis/assembly and cytokinesis are shared targets of selection in both haploid and diploid populations. However, even within these biological processes, there are both shared (e.g. *KRE6*) and ploidy-specific target genes, such as the haploid-specific *GAS1* and *KRE5*, and the diploid-specific *CTS1*. Furthermore, the most common haploid target of selection, *IRA1* (a negative regulator of Ras), was not mutated in any of our diploid populations.

**Table 1.** Common targets of selection in diploid populations

| GENE | MUTATIONS |  |  |  | GO Term (Biological Process)* |
| --- | --- | --- | --- | --- | --- |
|  | Total | Heterozygous | Homozygous | Mixed† |  |
| *CTS1* | 6 | 0 | 6 | 0 | Cell Separation After Cytokinesis |
| *ACE2* | 4 | 2 | 2 | 0 | Cell Separation After Cytokinesis |
| *BCK1* | 2 | 1 | 0 | 1 | Regulation of Cell Wall Organization |
| *BEM2* | 2 | 2 | 0 | 0 | Actin Cytoskeleton Organization/Cell Polarity/Neg Reg of Rho |
| *BPH1* | 2 | 2 | 0 | 0 | Cell Wall Organization |
| *BST1* | 2 | 2 | 0 | 0 | Vesicle Transport/GPI Anchoring |
| *EDE1* | 2 | 2 | 0 | 0 | Pos Reg of Cytokinesis/ER UPR/Endocytosis/Actin Organization |
| *FKS1* | 2 | 0 | 2 | 0 | 1-3 Beta Glucan Biosynthesis/Pos Reg of Endocytosis |
| *HPF1* | 2 | 1 | 1 | 0 | Cell Wall Organization |
| *KRE6* | 2 | 2 | 0 | 0 | 1-6 Beta Glucan Biosynthesis/Cell Wall Organization |
| *PDR1* | 2 | 2 | 0 | 0 | Pos Reg of Cell Drug Response/Pos Reg of Transcription from RNAPol2 Promoter |
| *PSE1* | 2 | 2 | 0 | 0 | Nuclear Export/Reg of Mitotic Nuclear Division |
| *SIZ1* | 2 | 2 | 0 | 0 | Chromosome Segregation/Neg Reg of Ubiquitination/Protein Sumoylation |
| *SNT2* | 2 | 2 | 0 | 0 | Histone Ubiquitination/Oxidative Stress Response |
| *UBP12* | 2 | 2 | 0 | 0 | Unknown/Thiol-specific Protease |
| *YDR090C* | 2 | 2 | 0 | 0 | Unknown/Integral Membrane Component |

\* GO Terms were manually curated from the *Saccharomyces* Genome Database (yeastgenome.org). GO-term analysis shows that these 16 genes are enriched for hydrolase activity acting on glycosyl bonds ( $p=0.00587$ )
† Mixed indicates populations where one clone contained a heterozygous mutation and the other clone contained a homozygous mutation

### Evolved alleles show varying degrees of dominance and ploidy-dependence

Given that cell wall mutants represent a diverse subset of beneficial mutations, including haploid-specific (*GAS1*, *KRE5*), diploid-specific (*CTS1*), and common (*KRE6*) beneficial mutational targets, we focused on these four targets of adaptation. We reconstructed these alleles as haploids, heterozygous diploids, and homozygous diploids and performed competitive fitness assays on all constructs to test whether the relative fitness effects of evolved alleles are contingent on the ploidy in which they arose (**Fig. 4**). Consistent with our classification of these mutations as putative drivers of adaptation, we found that each is beneficial in the background in which it arose. Mutations in the shared target of selection, *KRE6* are beneficial as haploids, heterozygous diploids, and homozygous diploids, though the benefit as a heterozygote is less than that of a homozygote, showing that it is partially dominant (coefficient of dominance, *h*=0.34). The diploid-specific target of selection, *CTS1*, is equally beneficial as a heterozygote or a homozygote (*h*≈1), however this mutation is neutral in a haploid background. Mutations in the haploid-specific targets, *GAS1* and *KRE5*, are beneficial in haploids and homozygous diploids. In the case of *KRE5*, the heterozygous diploid fitness is not statistically different from zero (*h*=0.10, **Fig. 4**). In the case of *GAS1*, the heterozygote is strongly deleterious.

**Fig. 4:**
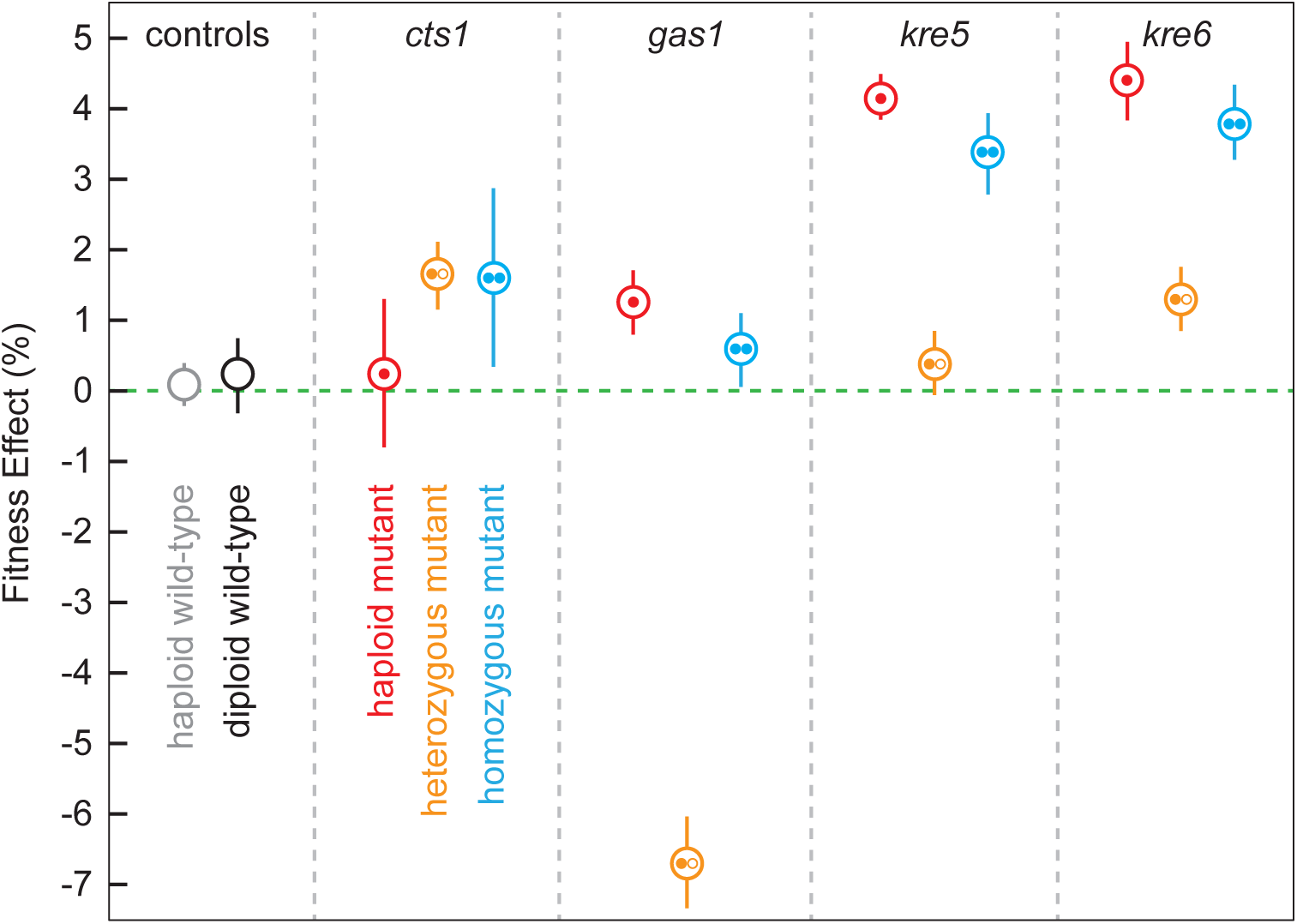
Evolved alleles show varying degrees of dominance and ploidy dependence. Average fitness effects of haploid (single filled circle in red), heterozygous diploid (one filled circle and one open circle in yellow) and homozygous diploid (two filled circles in blue) cell wall mutants are compared to wild-type haploids (light grey empty circle) and wild-type diploids (dark grey empty circle). The corresponding gene mutated in each is listed at the top. The *cts1* mutation is from the F05 population of our diploid data (**Table S1**). The *gas1*, *kre5*, *and kre6* mutations are from our haploid populations BYS2-D06, RMB2-B10, and RMS1-H08, respectively^14^. Each point is an average of seven replicates, or six replicates for the wild-type points. Error bars are the standard error of these averages.

### Gene conversion occurs more rapidly for *CTS1* than *ACE2*

Interestingly, although our reconstruction experiments show no significant fitness difference between a heterozygous and homozygous *CTS1* mutation, all six *CTS1* mutations are homozygous in the diploid populations. Another common diploid-specific target, *ACE2*, is observed as a homozygous mutant in two of the four populations in which it arose. Both *ACE2* and *CTS1* are located on Chr. XII. Two possible mechanisms could explain this observation: gene conversion or chromosome loss. To verify that our strains have two copies of Chr. XII, we measured read coverage in our sequencing data and found that coverage across Chr. XII was consistent with coverage across the rest of the genome in all samples containing a *CTS1* or *ACE2* mutant. We also performed tetrad dissections of *CTS1* mutant strains and found four-spore viable tetrads, which is unlikely if all or part of one copy of Chr. XII is missing. These homozygous mutations therefore likely arise from gene conversion events.

With this in mind, we investigated the dynamics of genome sequence evolution by performing whole-genome whole-population time-course sequencing on two populations, one with a homozygous *CTS1* mutation, and one with a homozygous *ACE2* mutation. In population A05, the *ace2* allele establishes around generation 900, rises to a frequency of ~0.5 by generation 1,100, and remains there until around generation 1,400, when it rises to a frequency of ~1, fixing in the population by generation 1,600 (**Fig. 5A**). In population F05, the *cts1* allele establishes around generation 700, rises to a frequency of ~0.5 around generation 800, and continues without pausing to reach a frequency of ~1 around generation 1,000 (**Fig. 5B**). In this case, the mutant *cts1* homozygote establishes before the heterozygote fixes. In addition to *CTS1* and *ACE2* mutations, we observe other mutations in these populations, including a few heterozygosities that were present in the ancestral strain. We can use the information from the dynamics of these additional mutations to inform our theory of gene conversion leading to homozygosity. The A05 population contains three non-*ACE2* variants, two of which sweep as heterozygotes. The third is an existing polymorphism in an *LTR* that was heterozygous in the ancestor and is approximately 387 kb away from *ACE2* on Chr. XII. When the *ace2* allele fixes in this population, the *LTR* also loses heterozygosity (**Fig. 5A**). The F05 population contains 11 non-*CTS1* mutations, including seven heterozygous mutations, two homozygous mutations, and two positions that were heterozygous in the ancestor, but lost heterozygosity during the evolution of this population. Among these is a mutation in *DUS4*, which sweeps to fixation with the *cts1* allele, and the previously described heterozygosity in an *LTR* which loses heterozygosity at the same time as *CTS1* (**Fig. 5B**). In this case, the *LTR* is approximately 84 kb away from *CTS1*.

**Fig. 5:**
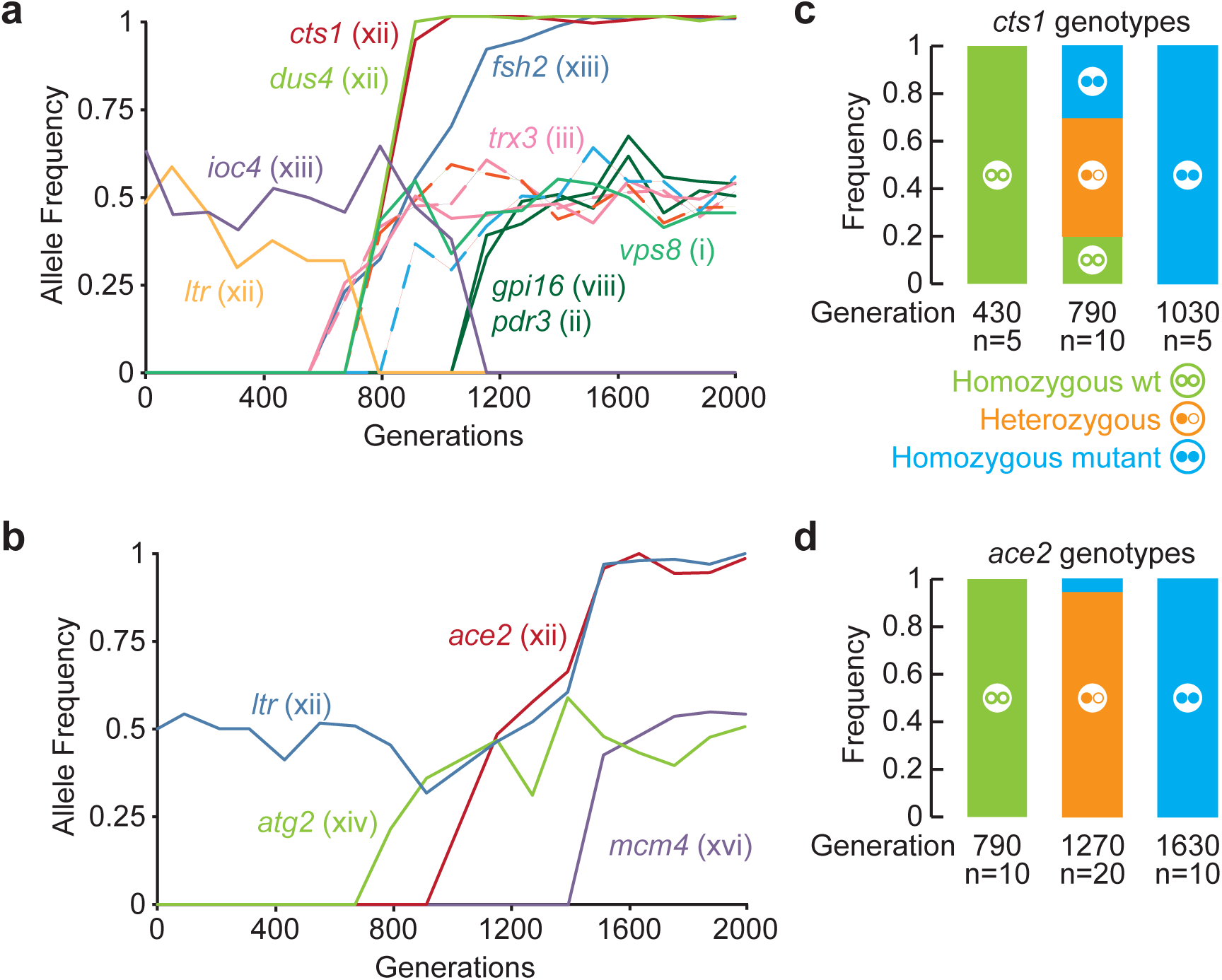
Dynamics of adaptation and gene conversion. **a,** Time-course sequencing of population F05 containing a *cts1* mutation shows multiple mutations fixing as a cohort with *cts1*. The *cts1* mutant fixes in the population quickly, without a pause around 0.5 signifying heterozygote fixation. Mutations are shown with the chromosomes on which they occur. Dashed lines represent intergenic mutations. **b,** Genotypes of *CTS1* at three time points are shown as homozygous wild type (two open circles), heterozygous (one filled and one open circle), and homozygous mutant (two filled circles). Homozygous mutants appear before homozygous wild type is eliminated. **c,** Time-course sequencing of population A05 containing an *ace2* mutation. There is a decrease in slope around an allele frequency of 0.5, suggesting heterozygote fixation before homozygous mutant establishment. Mutations are shown with the chromosomes on which they occur. **d,** Genotypes of *ACE2* at three time points are shown as homozygous wild type (two open circles), heterozygous (one filled and one open circle), and homozygous mutant (two filled circles). Homozygous mutants only appear after the homozygous wild type is eliminated by the sweep of the heterozygous mutant.

We hypothesize that the length of the pause of the *ace2* allele at a frequency of 0.5 is the waiting time between the heterozygote fixing in the population and the gene conversion event. To test this, we performed Sanger sequencing of clones from both populations before, during, and after the sweep, corresponding to allele frequencies of 0, ~0.5 and 1. For *ACE2*, 19/20 clones at an allele frequency of ~0.5 are heterozygous, consistent with a full sweep of the heterozygote before gene conversion (**Fig. 5C**). For *CTS1* at an allele frequency of ~0.5, we observe a mix of heterozygous, homozygous mutant, and homozygous ancestral clones, meaning that the heterozygote did not fix in the population before the gene conversion event, as we saw for the *ace2* allele in population A05 (**Fig. 5D**). This suggests that the *cts1* allele gene converts very quickly compared to the *ace2* allele.

While the F05 allele of *CTS1* is a nonsense mutation (likely to be a loss-of-function), the other five *CTS1* mutant alleles seen in our diploids are missense mutations (possibly alteration-of-function). Even so, all six show similar dynamics of adaptation, with similar rates of gene conversion (**Figure S1**). Based on this, we hypothesize that the difference in gene conversion rates between *CTS1* and *ACE2* mutations is likely not due to the effects of individual mutations, but rather is due to the location of the *CTS1* and *ACE2* genes themselves.

## DISCUSSION

We show that our diploid populations adapt more slowly than haploids because recessive beneficial mutations are not selectively accessible to diploids. We have used our power law and linear fits to predict future rates of adaptation for our evolving haploids and diploids, respectively. In our model, the rate of haploid adaptation starts off higher but decreases over time following the power law, while the diploid rate of adaptation is lower but constant, following a linear fit. Therefore, the model predicts that around generation 3,500, the haploid and diploid rates of adaptation are equal, but past this point, the diploid rate is faster than the haploid rate (**Fig. S1A**). Additionally, our model predicts that by generation 15,000, diploids will have achieved more total adaptation relative to their ancestor than haploids (**Fig. S1B**). The declining rate of adaptation in haploids may be due to the exhaustion of available beneficial recessive mutations.

Through our comparison of the mutations gained during haploid and diploid adaptation, we observe that both the mating pathway and the negative regulation of Ras were prominent targets of adaptation in haploids, but were not targets of selection in diploids. This makes sense for the mating pathway, as it is repressed in diploids (and therefore sterile mutations are not selectively advantageous). As to why negative regulation of Ras is not a target of adaptation in diploid populations, there are two possibilities. The first is that all or most of the spectrum of beneficial mutations that can be made to genes involved in the negative regulation of Ras are recessive in their fitness effects, and thus unlikely to pass through Haldane’s sieve. The other possibility is that the Ras pathway is regulated differently in haploids and diploids, such that some genes are not functionally redundant in haploids and diploids. Some yeast genes are known to have ploidy-specific regulation^5^. Interestingly, *CTS1* is included among these differentially regulated genes, increasing in expression with ploidy. This physiological difference between haploids and diploids may explain why mutations to *CTS1* are diploid-specific.

By examining our diploid evolved mutations, we found that Haldane’s sieve is effective at filtering out recessive beneficial mutations from evolving diploid populations, but it is not the only factor affecting the rate of adaptation and spectrum of beneficial mutations in diploids. We also see that genomic position is an important factor, largely due to varying rates of recombination throughout the genome. This is particularly visible on the right arm of Chr. XII, which contains the rDNA locus, a known recombination hotspot in yeast^22^. We find that homozygous mutations are rare (only 10% of diploid mutations) but are largely concentrated on the right arm of Chr. XII (**Fig. 3**), particularly in the *CTS1* and *ACE2* genes (**Fig. S2**). This implies that the ability for beneficial diploid mutations to become homozygous, and thus to escape from Haldane’s sieve will depend strongly on local rates of gene conversion. Our results here further validate the point made by Mandegar and Otto that mitotic recombination is an important factor in the spread of beneficial alleles in evolving asexual populations^23^.

We reconstructed evolved haploid-specific, diploid-specific, and shared alleles in isolation as haploids, heterozygous diploids, and homozygous diploids, and found that the degree of heterozygosity differs between the alleles. Our reconstructed allele of *CTS1* shows a surprisingly high degree of dominance (*h*≈1), yet all six of the *cts1* alleles present in our diploid populations rapidly gene converted and fixed as homozygotes. One possible explanation is that the homozygote does have an advantage over the heterozygote, but that we cannot detect this difference with flow cytometry-based fitness assays. One complication is that homozygous mutations in *CTS1* result in a cell aggregation phenotype. This aggregation phenotype has previously been shown to occur during yeast laboratory evolution^24–27^. Our evolved *CTS1*-mutant diploid populations and the reconstructed *cts1* haploid and homozygous diploid strains form aggregates, but the reconstructed heterozygous mutant does not (**Fig. S3**). It is possible that this phenotype complicates fluorescence-based fitness measurements of these strains.

Furthermore, from the dynamics of adaptation of all six *CTS1*-mutant populations (**Fig. 5A**, **Fig. S4**), we have shown that *CTS1* mutants become homozygous both very frequently and quickly. Additionally, from the dynamics of the F05 population containing our *CTS1* mutation of interest, we see a mutant *dus4* allele that travels to fixation with the *cts1* allele, and one ancestral heterozygosity in a long terminal repeat that loses heterozygosity at the same time (**Fig. 5A**). All three of these genetic loci are on Chr. XII downstream from the yeast rDNA locus. We propose that a single recombination event caused a loss of heterozygosity for all three of these loci, establishing the *cts1* homozygous mutant. Our dynamics of adaptation also show the loss of heterozygosity of another ancestral heterozygous long terminal repeat paired with the rise of the *ace2* allele to fixation, suggesting that a similar gene conversion may have established the *ace2* homozygote. In this same population, there may be another case of gene conversion where the loss of heterozygosity in *IOC4* co-occurs with the fixation of an *FSH2* mutant. Both of these loci are on Chr. XIII (**Fig. 5C**). Through these examples and the concentration of homozygous mutations and lack of heterozygous mutations on the right arm of Chr. XII, we also propose that when gene conversion events occur, they involve a single point of recombination leading to loss of heterozygosity from the point of origin through the telomere. This is similar to the types of lesions observed due to break-induced replication^28^.

While gene conversion allows us to explain the location-specific enrichment of homozygous mutations, these are a minority of our putative driver mutations (**Table 1**) and are therefore not sufficient to explain the fitness gains we see in diploids. There must also be some adaptive heterozygous mutations, which remain heterozygous. It is possible that some of our heterozygous putative driver mutations are maintained as heterozygotes because they are overdominant as suggested by Sellis *et al*.^17,18^. Overall, we have shown here that over the course of 4,000 generations of evolution, diploid populations adapt slower than haploid populations, and that diploid populations have a unique spectrum of beneficial mutations compared to haploid populations. In addition, the majority of adaptive diploid mutations are heterozygous and the prevalence of adaptive diploid homozygous mutations depends on the position of mutations in the genome. Collectively this work fills a gap in understanding how ploidy impacts adaptation, and provides empirical support for the hypothesis that diploid populations have restricted access to beneficial mutations.

## METHODS

### Strain Construction

The strains used in this experiment are derived from the base strain, yGIL432, a haploid yeast strain derived from the W303 background with genotype *MAT***a**, *ade2*-*1*, *CAN1*, *his3*-*11*, *leu2*-*3*,*112*, *trp1*-*1*, *URA3*, *bar1*Δ::*ADE2*, *hm*lαΔ::*LEU2*, *GPA1*::NatMX, *ura3*Δ::P*_FUS1_*-yEVenus. This strain was previously reported as DBY15105 ^25^. A nearly isogenic *MAT*α version of yGIL432 was constructed by introgressing the *MAT*α allele through three backcrosses into the yGIL432 background. This *MAT*α strain, yGIL646, was crossed to yGIL432 to generate the diploid strain yGIL672. Haploid *MAT*a evolved strains used in Figure 1 are previously described in Ref. 13, and their corresponding *MATa/a* heterozygous diploids are described in Ref. 20. For the reconstruction experiments, the haploid *cts1* mutation in the yGIL432 background was generated using Cas9 allele replacement^29^. Briefly we retargeted the guide RNA expressing plasmid (Addgene #43803) to *cts1* using the gRNA sequence 5′ TTCTTCAAAATCTCAACATA 3′. We co-transformed the gRNA plasmid (pGIL083), a constitutive Cas9 plasmid (Addgene #43802), and a plasmid (pGIL089) containing the *cts1* mutant allele into our *MAT*a strain, yGIL432. We screened for plasmid retention, PCR screened single colonies for the allele replacement, and cured the plasmids. Allele replacements of *gas1*, *kre5*, and *kre6* in the yGIL432 were obtained from Matt Remillard (Princeton University), who constructed them using alleles from our haploid populations BYS2-D06, RMB2-B10, and RMS1-H08, respectively. Each of the haploid *MAT*a strains was crossed to yGIL646 to generate heterozygous diploids. The heterozygous diploids were then sporulated, tetrads were dissected, and haploid spores were mating-type tested and genotyped. Appropriate haploid spores were then crossed to each other to create homozygous diploid strains for each mutation.

### Long-Term Evolution

To set up the long-term evolution experiment, a single clone yGIL672 was grown to saturation in rich-glucose YPD medium, was diluted 1:10,000, and was used to seed 48 replicate populations in a single 96-well plate. This initial plate was duplicated and then frozen down for future use. The cultures were evolved through 4,000 generations (400 cycles) of growth and dilution in YPD at 30°C. Every 24 hours, the populations were diluted 1:1024 by serial diluting 1:32 (4 μl into 130 μl) x 1:32 (4 μl into 130 μl) into new medium containing ampicillin (100 mg/L) and tetracycline (25 mg/L). All dilutions were performed using the Biomek Liquid Handler equipped with a Pod96. Approximately every 50 generations, plates were mixed with 50 μl of 60% glycerol and archived at −80°C.

### Competitive Fitness Assays

Flow cytometry-based competitive fitness assays were performed essentially as described previously^13,20,25^ Briefly, experimental and reference strains were grown to saturation in separate 96-well plates. If the strains to be tested were coming from the freezer, the strains were passaged once by diluting 1:1024 to re-acclimate the strains to the appropriate medium. Experimental and reference strains were mixed 50:50 using the Biomek Liquid handler and were propagated for 40 generations under identical conditions to the original evolution experiment. In this project we used two reference strains: diploid *MAT***a**/**a**, and diploid *MAT***a**/α. These reference strains are derived from the yGIL432 base strain, but contain a constitutive ymCitrine integrated at the *ura3* locus.

We used previously collected competitive fitness data from evolved *MAT***a** haploid strains^13^. We performed a competitive fitness assay as described below on previously constructed heterozygous *MAT***a**/**a** diploid strains^20^. To perform this assay, we constructed a *MAT***a**/**a** diploid version of the fluorescently-labeled reference strain using a plasmid containing *LEU2* under a *MAT***a**-specific promoter to select for gene conversion at the mating type locus. These data were normalized to their respective references and compiled into Figure 1.

### Whole-Genome Sequencing

For sequencing clones, we struck to single colonies on YPD and picked two colonies from each population to sequence. These individuals were grown to saturation in liquid media and total genomic DNA was isolated for each sample. For whole-genome whole-population time-course sequencing, we thawed each population at 18 time points from generation 0 to 2,000, and transferred 10 μl into 5 ml of YPD. We made genomic DNA preparations as above.

We followed a modified version of the Nextera sequencing library preparation protocol^30^, modified further as described in Ref. 20. We used the Nextera sequencing library preparation kit and protocol to isolate total genomic DNA and add the unique Nextera library barcodes to all 48 samples. We measured the concentration of DNA in each sample using a NanoDrop spectrophotometer and confirmed these values using a Qubit fluorometer. We equalized the DNA concentration of each sample via dilution and mixed all 48 samples into a single pool. We used a BioAnalyzer High-Sensitivity DNA Chip (BioAnalyzer 2100, Agilent) to confirm that the pool contained the appropriate length DNA fragments, and performed gel extraction on the pool to remove short fragments. The pool was run on an Illumina HiSeq 2500 sequencer with 157 nucleotide single-end reads by the Sequencing Core Facility within the Lewis-Sigler Institute for Integrative Genomics at Princeton University. After an initial sequencing run provided us with the number of reads from each sample, we remixed the pool to better represent underrepresented samples and it was resequenced.

### Sequencing Analysis Pipeline

The raw sequencing information was first merged from 3 lanes of sequencing via concatentation. This single file was split into 48 files by the barcodes corresponding to each sample using a custom Perl script (barcode_splitter.py) from L. Parsons (Princeton University). Each sample was aligned to the same customized W303 genome based on the S288C genome available on SGD using the Burrows-Wheeler Aligner (BWA, Version 0.7.12), using default parameters except “Disallow an indel within INT bp towards the ends” set to 0 and “Gap open penalty” set to 5, creating both a .bam and .bai file for each. Variants were called using the FreeBayes variant caller (Version 0.9.21-24-g840b412) and merged together into a single spreadsheet. Variants that existed only in paired clones (448 mutations total) were annotated manually using the Integrated Genome Viewer (IGV) and false calls were removed. This fully annotated table (containing 383 mutations) is available in Table S1.

For whole-genome whole-population time-course sequencing, we used the same Nextera sequencing library preparation protocol as described above, with the following changes: Instead of two clones from each of 24 populations, we isolated genomic DNA from whole populations at 18 time points for two populations for a total of 36 samples. During the sequencing analysis, after splitting the reads based on the Nextera barcodes, we used a script (clipper.sh) to remove any Nextera adapter sequences introduced by sequencing short fragments. We used a previously-described set of scripts (allele_counts.pl and composite_scores.pl) to call real mutations^13^.

### Reconstruction Experiments

We used three previously identified mutations, which were identified in evolved haploids (mutations to *gas1*, *kre5*, and *kre6*), and one mutation identified in a diploid population (mutation to *cts1*). We used CRISPR/Cas9 genome editing to reconstruct our evolved *cts1* mutation in our ancestral haploid background. The other three mutations were previously reconstructed as single mutants in our haploid background by Matt Remillard (Princeton University), who generously shared these resources with us. These four haploid *MAT***a** strains were crossed to our haploid *MAT*α ancestor to construct heterozygous diploids for each mutation. These heterozygous diploids were sporulated, tetrads were dissected, and haploid spores were mating-type tested and Sanger-sequenced to determine which spores contained the mutation but were of opposite mating type. These haploid spores were then crossed to create diploids homozygous for each mutation. Fitness of each of these twelve total samples was measured via competitive fitness assays across seven replicates along with control haploids and diploids with no mutations, all against the appropriate fluorescent ancestor strains.

## ACKNOWLEDGEMENTS

We thank Matthew Remillard (Princeton University) for providing strains. We thank Alex Nguyen and Michael Desai (Harvard University) for providing the plasmid with mating-type-specific selectable markers. We thank Anna Selmecki, Sean Buskirk, Katie Fisher, and Ryan Vignogna for their comments on the manuscript. This work was supported by the Charles E. Kaufman Foundation of The Pittsburgh Foundation.

## FIGURE LEGENDS

**Fig. S1:**
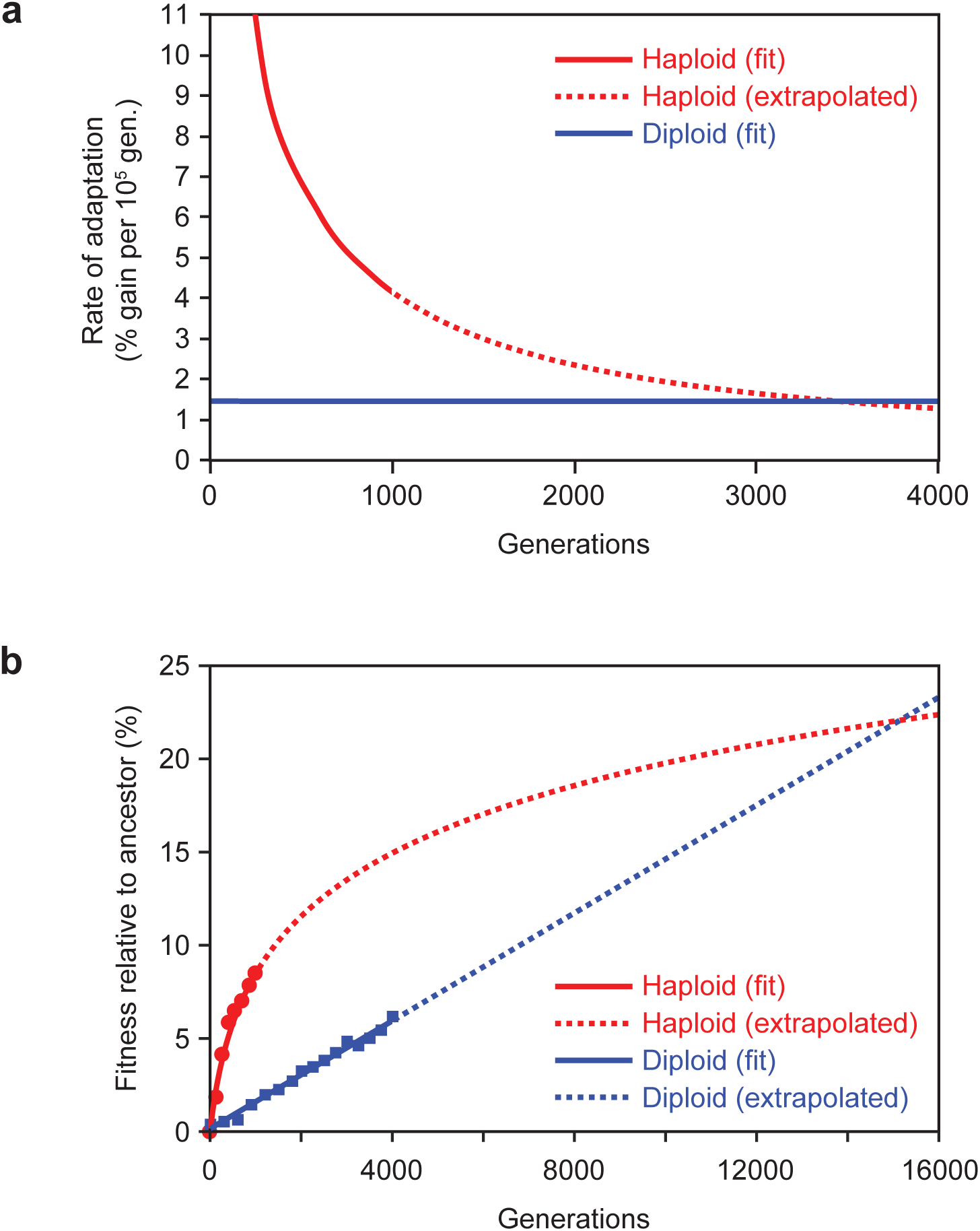
Predicting rate of adaptation and total amount of adaptation over time using power law and linear fit data. **a,** The average rate of adaptation for haploids (red) and diploids (blue). The haploid rate decreases over time, while the diploid rate remains constant. The red dotted line is the extrapolation of the haploid rate of adaptation based on the power law. **b,** Average adaptation relative to ancestor for haploids (red), and diploids (blue), with predictions out to 16,000 generations using power law for haploid data (red dotted line) and linear fit for diploid data (blue dotted line).

**Fig. S2:**
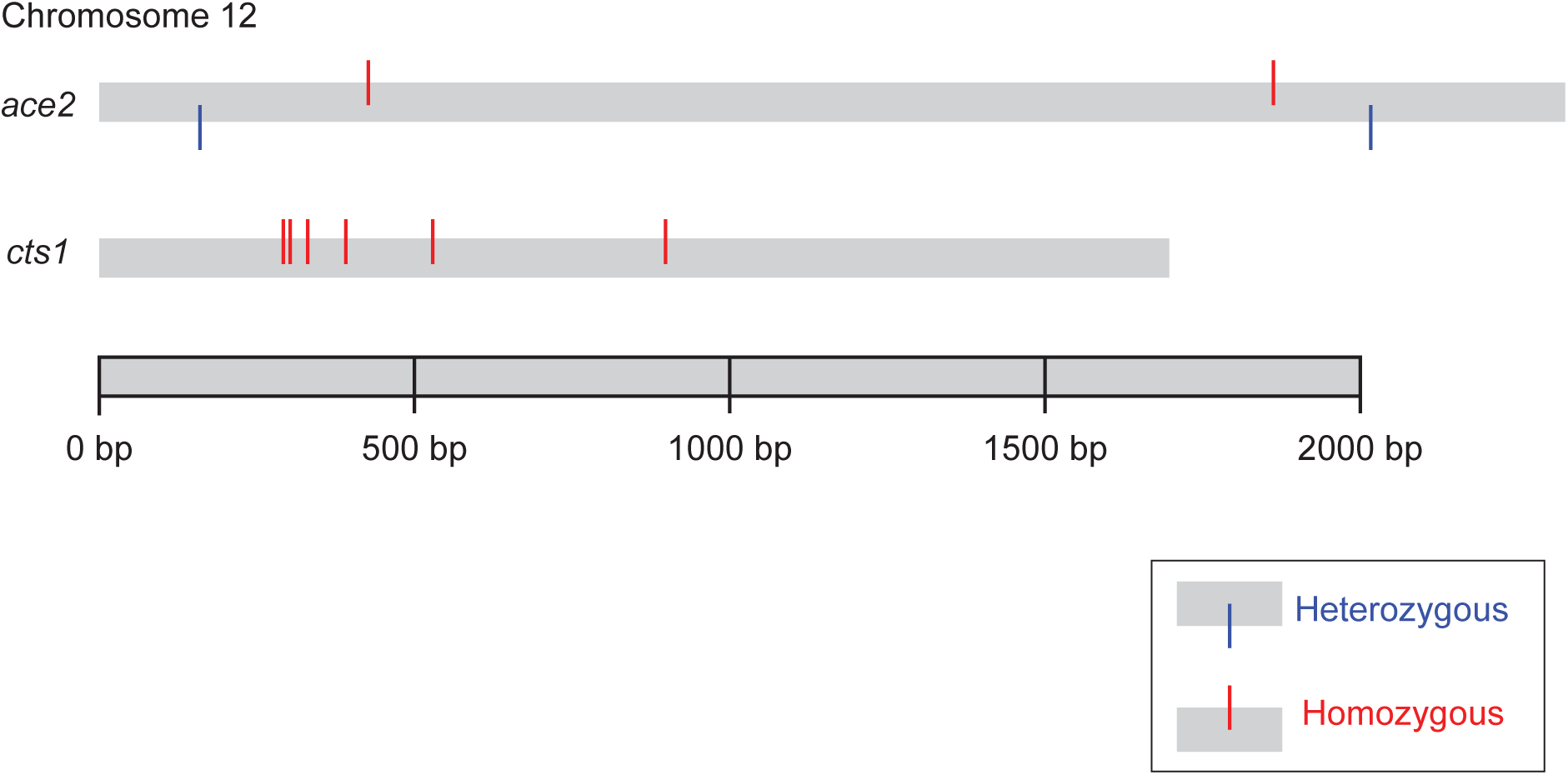
Distribution of mutations to *ACE2* and *CTS1* in diploids evolved for 2,000 generations. The *ACE2* locus on Chr. XII is the site of two homozygous and two heterozygous mutations. The *CTS1* locus on Chr. XII is the site of six homozygous mutations. Homozygous mutations are represented by red lines sticking out of the top of chromosomes. Heterozygous mutations are represented by blue lines sticking out of the bottom of chromosomes.

**Fig. S3:**
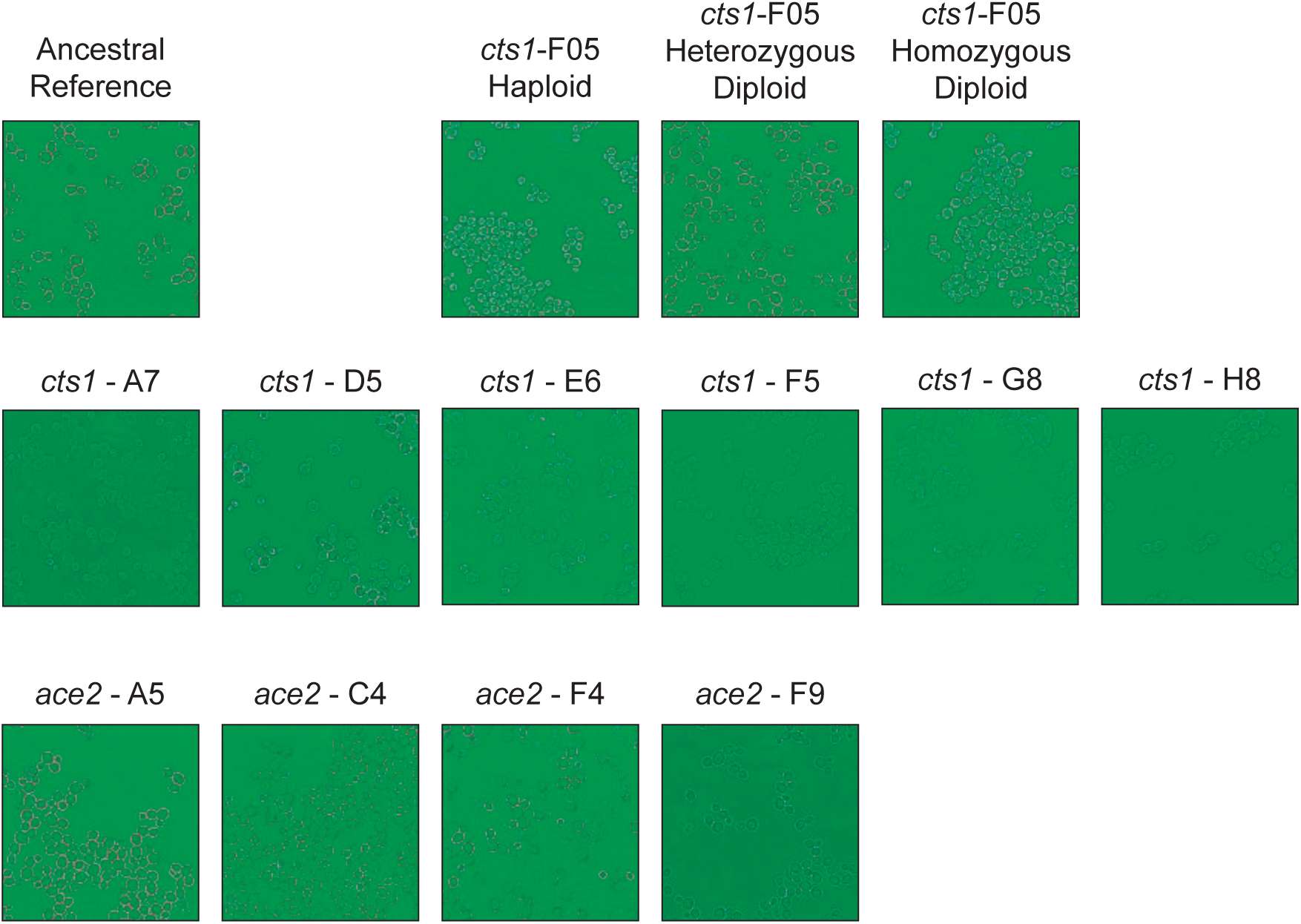
Aggregation phenotypes occur in some cell wall mutants. The reference is the common ancestor of all strains shown here. The top row shows reconstructed single-mutant strains for the F05-*cts1* allele. The middle row shows the six evolved populations which contain *cts1* mutations. The bottom row shows the four evolved populations which contain *ace2* mutations. Each image was taken at 400x magnification.

**Fig. S4:**
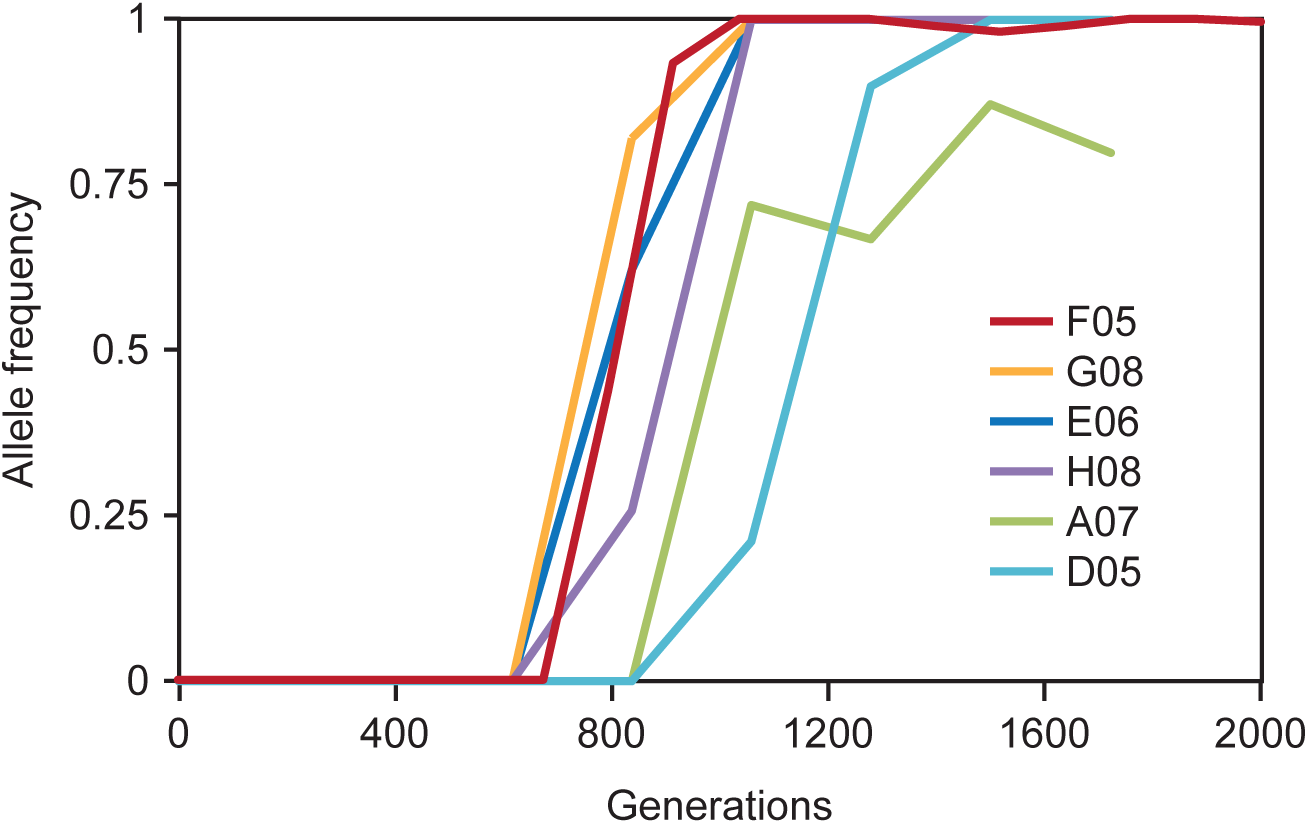
All six *cts1* mutants show similar dynamics. Sanger sequencing of five additional *cts1* mutants shows that the dynamics of adaptation are similar between *cts1* mutants. The six total populations are listed on the right.

## REFERENCES

1 Hummer, K. E., Nathewet, P. & Yanagi, T. Decaploidy in Fragaria iturupensis (Rosaceae). American journal of botany 96, 713–716, doi:10.3732/ajb.0800285 (2009).

2 Goodenough, U. & Heitman, J. Origins of eukaryotic sexual reproduction. Cold Spring Harbor perspectives in biology 6, doi:10.1101/cshperspect.a016154 (2014).

3 Orr, H. A. & Otto, S. P. Does diploidy increase the rate of adaptation? Genetics 136, 1475–1480 (1994).

4 Haldane, J. B. S. A mathematical theory of natural and artificial selection, part V: selection and mutation. Pros. Cambridge Philos. Soc. 23, 838–844 (1927).

5 Galitski, T., Saldanha, A. J., Styles, C. A., Lander, E. S. & Fink, G. R. Ploidy regulation of gene expression. Science (New York, N.Y.) 285, 251–254 (1999).

6 Paquin, C. & Adams, J. Frequency of fixation of adaptive mutations is higher in evolving diploid than haploid yeast populations. Nature 302, 495–500 (1983).

7 Anderson, J. B., Sirjusingh, C. & Ricker, N. Haploidy, diploidy and evolution of antifungal drug resistance in Saccharomyces cerevisiae. Genetics 168, 1915–1923, doi:10.1534/genetics.104.033266 (2004).

8 Gerstein, A. C., Cleathero, L. A., Mandegar, M. A. & Otto, S. P. Haploids adapt faster than diploids across a range of environments. Journal of evolutionary biology 24, 531–540, doi:10.1111/j.1420-9101.2010.02188.x (2011).

9 Selmecki, A. M. et al. Polyploidy can drive rapid adaptation in yeast. Nature 519, 349–352, doi:10.1038/nature14187 (2015).

10 Zeyl, C., Vanderford, T. & Carter, M. An evolutionary advantage of haploidy in large yeast populations. Science (New York, N.Y.) 299, 555–558, doi:10.1126/science.1078417 (2003).

11 Kryazhimskiy, S., Rice, D. P., Jerison, E. R. & Desai, M. M. Global epistasis makes adaptation predictable despite sequence-level stochasticity. Science (New York, N.Y.) 344, 1519–1522, doi:10.1126/science.1250939 (2014).

12 Kvitek, D. J. & Sherlock, G. Whole genome, whole population sequencing reveals that loss of signaling networks is the major adaptive strategy in a constant environment. PLoS genetics 9, e1003972, doi:10.1371/journal.pgen.1003972 (2013).

13 Lang, G. I. et al. Pervasive genetic hitchhiking and clonal interference in forty evolving yeast populations. Nature 500, 571–574, doi:10.1038/nature12344 (2013).

14 Tenaillon, O. et al. The molecular diversity of adaptive convergence. Science (New York, N.Y.) 335, 457–461, doi:10.1126/science.1212986 (2012).

15 Scott, A. L., Richmond, P. A., Dowell, R. & Selmecki, A. M. The influence of polyploidy on the evolution of yeast grown in a sub-optimal carbon source. Mol Biol Evol, doi: 10.1093/molbev/msx205 (2017).

16 Gerstein, A. C., Kuzmin, A. & Otto, S. P. Loss-of-heterozygosity facilitates passage through Haldane’s sieve for Saccharomyces cerevisiae undergoing adaptation. Nature communications 5, 3819, doi:10.1038/ncomms4819 (2014).

17 Sellis, D., Callahan, B. J., Petrov, D. A. & Messer, P. W. Heterozygote advantage as a natural consequence of adaptation in diploids. Proceedings of the National Academy of Sciences of the United States of America 108, 20666–20671, doi:10.1073/pnas.1114573108 (2011).

18 Sellis, D., Kvitek, D. J., Dunn, B., Sherlock, G. & Petrov, D. A. Heterozygote Advantage Is a Common Outcome of Adaptation in Saccharomyces cerevisiae. Genetics 203, 1401–1413, doi:10.1534/genetics.115.185165 (2016).

19 Deutschbauer, A. M. et al. Mechanisms of haploinsufficiency revealed by genome-wide profiling in yeast. Genetics 169, 1915–1925, doi:10.1534/genetics.104.036871 (2005).

20 Buskirk, S. W., Peace, R. E. & Lang, G. I. Hitchhiking and epistasis give rise to cohort dynamics in adapting populations. Proceedings of the National Academy of Sciences of the United States of America, doi:10.1073/pnas.1702314114 (2017).

21 Wiser, M. J., Ribeck, N. & Lenski, R. E. Long-term dynamics of adaptation in asexual populations. Science (New York, N.Y.) 342, 1364–1367, doi:10.1126/science.1243357 (2013).

22 Keil, R. L. & Roeder, G. S. Cis-acting, recombination-stimulating activity in a fragment of the ribosomal DNA of S. cerevisiae. Cell 39, 377–386 (1984).

23 Mandegar, M. A. & Otto, S. P. Mitotic recombination counteracts the benefits of genetic segregation. Proceedings. Biological sciences 274, 1301–1307, doi:10.1098/rspb.2007.0056 (2007).

24 Koschwanez, J. H., Foster, K. R. & Murray, A. W. Sucrose utilization in budding yeast as a model for the origin of undifferentiated multicellularity. PLoS biology 9, e1001122, doi:10.1371/journal.pbio.1001122 (2011).

25 Lang, G. I., Botstein, D. & Desai, M. M. Genetic variation and the fate of beneficial mutations in asexual populations. Genetics 188, 647–661, doi:10.1534/genetics.111.128942 (2011).

26 Oud, B. et al. Genome duplication and mutations in ACE2 cause multicellular, fast-sedimenting phenotypes in evolved Saccharomyces cerevisiae. Proceedings of the National Academy of Sciences of the United States of America 110, E4223–4231, doi:10.1073/pnas.1305949110 (2013).

27 Ratcliff, W. C., Denison, R. F., Borrello, M. & Travisano, M. Experimental evolution of multicellularity. Proceedings of the National Academy of Sciences of the United States of America 109, 1595–1600, doi:10.1073/pnas.1115323109 (2012).

28 Kraus, E., Leung, W. Y. & Haber, J. E. Break-induced replication: a review and an example in budding yeast. Proceedings of the National Academy of Sciences of the United States of America 98, 8255–8262, doi:10.1073/pnas.151008198 (2001).

29 Mans, R. et al. CRISPR/Cas9: a molecular Swiss army knife for simultaneous introduction of multiple genetic modifications in Saccharomyces cerevisiae. FEMS yeast research 15, doi:10.1093/femsyr/fov004 (2015).

30 Baym, M. et al. Inexpensive multiplexed library preparation for megabase-sized genomes. PloS one 10, e0128036, doi:10.1371/journal.pone.0128036 (2015).

